# From single scenes to extended scenarios: the role of the ventromedial prefrontal cortex in the construction of imagery-rich events

**DOI:** 10.1101/2025.05.03.652053

**Authors:** Julia Taube, Pitshaporn Leelaarporn, Maren Bilzer, Ruediger Stirnberg, Yilmaz Sagik, Cornelia McCormick

## Abstract

Mental events are fundamental to daily cognition, including the recollection of past experiences, the anticipation of future scenarios, and engagement in imaginative, fictitious thought. Typically, these temporally extended mental events unfold within coherent spatial contexts, rich in naturalistic scenes and objects. However, there remains a significant gap in understanding how these events are represented in the brain. This study aimed to investigate the neural patterns involved in the construction of temporally extended mental events. Using ultra-high field functional magnetic resonance imaging, we examined brain regions previously implicated in this cognitive process, including the ventromedial prefrontal cortex (vmPFC), hippocampus, and posterior neocortex. We employed a novel experimental paradigm in which participants engaged in three forms of mental imagery: single objects (e.g., “a black espresso”), single scenes (e.g., “a busy café”), and extended scenarios (e.g., “meeting a friend for coffee”). We identified a shared neural network - comprising the vmPFC, hippocampus, and posterior neocortex – engaged across all forms of mental imagery. However, we observed a hierarchical organization in their contributions: the posterior neocortex supported the construction of objects, scenes, and scenarios, while the hippocampus primarily contributed to scenes and scenarios. The vmPFC exhibited a stepwise increase in activation, peaking during scenario construction. These findings suggest that the construction of mental events emerges from the close interaction of perceptual details provided by the posterior neocortex, spatial coherence from the hippocampus, and the integration of those elements into a coherent, temporally extended mental event by the vmPFC – the “movies” of the mind.

## Introduction

Mental events are fundamental to everyday cognition, enabling the recollection of autobiographical events, the envisioning of our future selves, and the creation of imagined scenarios. For most individuals, these movie-like mental events unfold within a coherent spatial framework, where naturalistic scenes and objects serve as key features. While extensive research has linked the construction of naturalistic scenes to the hippocampus [1–6], the specific contributions of the ventromedial prefrontal cortex (vmPFC), and posterior neocortex in constructing these temporally extended mental movies remain less understood. To distinguish terminology, we use “mental events” as a broader term for all types of mental activity including autobiographical memory recall, future thinking, etc.

We have recently suggested that the vmPFC plays a pivotal role that goes beyond mentally constructing individual objects and scenes [7]. Within this framework the vmPFC is critical for initiating and elaborating temporally extended, imagery-rich mental events – hereafter referred to as “scenarios”. This view emerged from a detailed examination of the cognitive effects of bilateral hippocampal damage, and bilateral vmPFC lesions First, we noticed that the spontaneous initiation of endogenous mental events (e.g., mind-wandering episodes) appeared reduced in vmPFC-damaged patients [8], and in the presence of hippocampal lesions [9–11]. Specifically, these patients exhibited lower rates of mind-wandering across various tasks and reported less frequent daydreaming compared to both healthy individuals and brain-damaged controls. Interestingly, vmPFC damage was associated with a decrease in off-task thoughts related to the future, while thoughts about the present were promoted. Second, while hippocampal-damaged patients often reported spatially fragmented individual scenes and autobiographical episodes with fewer spatial elements [12–14], vmPFC-damaged patients seem to be able to recall and describe coherent individual scenes and only showed impairments when constructing extended and complex scenarios [15]. Together, these findings suggest that while the vmPFC is not directly responsible for the visuo-spatial or mnemonic aspects of scene construction, it may be essential for the temporal unfolding of a dynamic mental scenario composed of scene imagery [16]. Consistent with this view, a meta-analysis found that the vmPFC was more strongly associated with the term “event” than with “scene” [17].

We propose that the vmPFC initiates the activation of specific scenarios and conveys this information to the hippocampus, which constructs individual scene snapshots from the broader scenario [18]. The vmPFC then engages in iterative feedback loops with the hippocampus and posterior neocortex, progressively integrating successive scenes and perceptual details into a temporally unfolding mental scenario.

Several magnetoencephalography (MEG) studies provide strong support for this model, demonstrating that the vmPFC plays a pivotal role in driving hippocampal scene construction processes [19–22]. McCormick et al. (2020) demonstrated that the vmPFC responds faster than the hippocampus during the initiation of autobiographical memory retrieval driving hippocampal evoked responses. Monk et al. (2020) further elucidated this interaction by showing that the vmPFC drives activity in the hippocampus during scene construction. Monk et al. (2021) expanded on these findings by demonstrating that the same interaction between the vmPFC and hippocampus occurs during tasks where scenes are integrated into the unfolding of a broader scenario.

Scene Construction Theory posits that the hippocampus plays a central role in generating spatially coherent mental representations by integrating disparate elements into a unified scene. This process involves the anterior hippocampus, which is thought to support the generation of novel spatial contexts and the binding of multimodal information, such as spatial, temporal, and sensory details, into a cohesive framework [23–26]. Previous research specifically connects scene construction to the hippocampus, with evidence highlighting the significant involvement of its anterior segment [3–5,7,12,27].

Thus, the current fMRI study investigates the precise contributions of the vmPFC, hippocampus, and posterior neocortex to the construction of naturalistic temporally extended mental imagery-rich scenarios. Using 7T MRI, we were able to achieve unprecedented spatial resolution and signal-to-noise ratio, allowing for more precise mapping of neural activity within our key regions. This level of detail is particularly crucial for understanding scenario construction, as it enables (1) the identification of subtle activation differences that may be obscured at lower field strengths and (2) the detection of fine-grained connectivity patterns. By leveraging these advantages, our study provides novel insights into the hierarchical organization of brain regions involved in constructing mental imagery-rich events.

Our hypotheses were as follows: (1) We expect the vmPFC to exhibit greater activation during scenario construction compared to scene and object construction. (2) We expect the hippocampus, particularly the anterior segment, to be more engaged during scenario and scene construction than during object construction. (3) We expect the posterior neocortex to be involved in all three types of imagery. This study has been pre- registered at https://doi.org/10.17605/OSF.IO/C46YR.

## Materials and methods

### Participants

Twenty-two healthy, right-handed participants were initially enrolled in the study after providing oral and written informed consent between 05.12.2022 and 30.10.2023.

Three participants were excluded due to suboptimal performance in the classification task during scanning (i.e., they performed in the 4^th^ percentile in at least one imagery category). The final sample comprised of nineteen participants (11 female, 8 male) with no history of neurological or psychiatric disorders and a mean age of 27.89 ± 3.67 years. For our experiment, it was crucial that participants reported a typical ability to visualize mental content. To assess this, we administered a German version of the Vividness of Visual Imagery Questionnaire (VVIQ), a 16-item scale that asks respondents to internally visualize four different scenarios and rate the vividness of their mental images on a 5-point Likert scale, yielding a total subjective vividness score [28]. Our exclusion criterion was a VVIQ score below 32, to exclude individuals with aphantasia [29]. No participants were excluded based on their VVIQ score. The mean score of our participants was 58.63 ± 10.39, indicating that they showed typical imagery ability.

To control for depressive symptoms, we included the German Beck Depression Inventory V (BDI-V), which assesses depressive symptoms across 21 items, each rated on a 6-point Likert scale, with a cut off score of 35 [30]. Mean scores were 20.78 ± 8.51. This study was reviewed and approved by the local ethics board of the University Hospital Bonn, Germany (Proposal 383/22). The study has been pre-registered to osf.io (https://doi.org/10.17605/OSF.IO/C46YR).

### Stimuli selection

The stimuli comprised four types of closely matched words representing an object, a scene, a scenario, or a non-word (a control task not involving visual construction). Most of the stimuli were selected from a previous study on imageability of word pairs and translated into German [27]. For the scenario condition, we incorporated novel stimuli. Before the main experiment, an independent sample of 23 healthy participants rated the type of imagery for 165 original words in an online questionnaire to confirm that each word triggered the intended type of imagery (object, scene, or scenario). Based on these ratings, we implemented a selection process based on discrimination accuracy to ensure optimal stimulus quality.

For the fMRI experiment, we selected only those stimuli that demonstrated the highest categorical discrimination (>80% accuracy), maximizing the reliability of neural responses during scanning. Stimuli with moderate discrimination rates (>60% accuracy) were allocated to the eye-tracking experiment. The remaining stimuli, which showed the lowest discrimination values while still reaching accuracy levels above chance, served as lures in the source memory task after scanning.

Based on accuracy values, stimuli in the fMRI experiment reached 87% (SD = 5%) for objects, 86% (SD = 4%) for scenes, and 88% (SD = 5%) for scenarios. In the eye-tracking experiment, accuracy was 73% (SD = 5%) for objects, 72% (SD = 6%) for scenes, and 73% (SD = 6%) for scenarios. For the source memory task, accuracy was 75% (SD = 4%) for objects, 63% (SD = 5%) for scenes, and 73% (SD = 6%) for scenarios. Stimuli of each imagery type (i.e., object, scene, scenario, and non-words) were matched for word length. In the eye-tracking task, average word lengths were 10.95 (SD = 5.60) for objects, 13.00 (SD = 7) for scenes, , and 11.00 (SD = 4.21) for scenarios. In the fMRI task, means were 11.95 (SD = 4.97) for objects, 12.45 (SD = 6.14) for scenes, 14.25 (SD = 6.02) for scenarios, and 11.65 (SD = 4.15) for non-words. In the source memory task, means were 13.65 (SD = 5.78) for objects, 13.40 (SD = 8.02) for scenes, and 11.45 (SD = 4.73) for scenarios. Word length did not differ significantly between the conditions (*p’s* > .05).

### General experimental procedure

Participants first received a detailed introduction, including task explanations and examples of all imagery types. After demonstrating understanding of the differences between object, scene, and scenario imagery, they completed a computer-based practice session. Once they answered correctly and had no further questions, the eye-tracking session was conducted, followed by the fMRI experiment. After the MRI session, participants completed a source memory task.

### Imagery task during eye-tracking and fMRI

During the eye-tracking and fMRI experiment, participants were asked to engage in three types of mental imagery (object, scene, or scenario construction) and a control task. During imagery trials, participants internally constructed and maintained a vivid mental image of the cued object, scene, or scenario while avoiding reliance on autobiographical memory.

For object imagery (e.g., a black espresso), participants were instructed to visualize a single, detailed version of the object in isolation, akin to a catalogue photo (e.g., a shiny, black espresso in a small red cup with a fish-scale pattern and a green handle). For scene imagery (e.g., a busy café), they constructed a spatially coherent, static image integrating multiple elements, resembling a postcard-like view (e.g., a street café in a picturesque Italian village with guests sitting at round tables under blue umbrellas, waiters carrying trays with cups and plates and colourful houses in the background).

To distinguish scenes from scenarios, participants were instructed to avoid motion in the scene condition. While motion implied, such as waiters serving guests, participants were instructed to mentally freeze the image and focus on the scene as a static, cohesive whole. In contrast, during scenario imagery (e.g., meeting a friend for coffee), participants were instructed to construct a mental simulation of a dynamic scenario that unfolds over time before their mind’s eye. For example, meeting a friend for coffee could involve greeting them at a busy train station, walking to a café, and catching up over coffee. Scenario construction differed from object and scene imagery by incorporating temporally extended scenarios with multiple scenes and objects. While all three conditions required mental imagery, they varied in their demands on scene construction and temporal complexity.

Extensive training was given to ensure that participants understood these instructions correctly and were able to differentiate between the three imagery conditions. Moreover, the eye-tracking task served as an extended practice session to ensure that participants fully understood the imagery tasks perfectly before entering the MRI scanner.

During the control task (only in the fMRI experiment), participants were required to count the number of characters in meaningless letter strings, matched in length to the experimental cues. The non-word counting task was chosen as a control condition because it requires basic cognitive engagement (e.g., attention and working memory) without involving mental imagery or constructive processes. This control condition allowed us to observe differences in brain activation attributable to imagery construction rather than general cognitive demands.

### Eye-tracking experiment

Eye movements were recorded using a video-based eye tracker (EyeLink 1000, SR Research) at a sampling rate of 2000 Hz. The system tracked participants’ eyes while their head position was stabilized using a chin rest positioned 64 cm from the 47 cm wide screen display and 58 cm from the desktop-mounted camera. Before the experiment started, a 9-point calibration and validation procedure were performed to ensure accurate tracking (average error < 0.5° of visual angle).

During the eye-tracking task, participants were shown 60 pseudo-randomized stimuli across one session, containing 20 scenario, 20 scene, and 20 object stimuli. Each trial began with a one-second fixation cross, followed by a one-second stimulus and a five-second blank display. During these five seconds, they were instructed to visualize the stimuli as detailed as possible in their mind’s eye without relying on past experiences. The stimulus was presented on a grey background to avoid afterimages in the blank display. This was followed by a (1) classification task, where participants categorized their imagined content as an object, scene, or scenario, and a (2) vividness rating, where they rated their perceived vividness on a 4-point Likert scale, ranging from very vague to very vivid. Participants responded in a self-paced manner (up to a maximum of three seconds). The experiment was run using SR Research Experiment Builder v. 1.1.

### MRI experiment

#### MR image acquisition

Structural and functional MRI data were acquired using a MAGNETOM 7 T Plus ultra-high field scanner (Siemens Healthineers, Erlangen, Germany).

As previously described [26], a whole-brain T1-weighted multi-echo MP-RAGE scan with 0.6 mm isotropic resolution was acquired using a custom sequence optimized for scanning efficiency and minimal geometric distortions [31,32]. The scan parameters were TI = 1.1 s, TR = 2.5 s, TEs = 1.84/3.55/5.26/6.92 ms, FA = 7°, TA = 7:12, readout pixel bandwidth: 970 Hz, matrix size: 428 x 364 x 256, elliptical sampling, sagittal slice orientation, CAIPIRINHA 1 x 2_z1_ parallel imaging with on-line 2D GRAPPA reconstruction, and a turbofactor of 218. The four echo time images were combined into a single high-SNR image using a root-mean-squares combination.

For functional imaging, a custom interleaved multishot 3D echo planar imaging (EPI) sequence was used with the following parameters: TE = 21.6 ms, TR_vol_ = 3.4 s, FA = 15°, 6/8 partial Fourier sampling, oblique-axial slice orientation along the anterior-posterior commissure line, readout pixel bandwidth: 1136 Hz, matrix size: 220 x 220 x 140. This sequence achieved a high spatial resolution of 0.9 mm isotropic at 7 T with sufficient SNR and a BOLD-optimal TE by combining several features: (A) Skipped-CAIPI 3.1 x 7_z2_ sampling [33] with on-line 2D GRAPPA reconstruction, (B) one externally acquired phase correction scan per volume, (C) variable echo train lengths with a semi-elliptical k-space mask [34], and (D) rapid slab-selective binomial-121 water excitation. A 3-min fMRI practice run was followed by two main functional sessions (∼15 min each). A standard 3 mm isotropic two-echo gradient-echo field-mapping scan was acquired in 35 s. A maximum of 264 imaging volumes were acquired per session, excluding the first five images to avoid non-steady-state signals.

### fMRI visual imagery task

Directly prior to the scanning procedure, participants underwent a short practice to get accustomed to the response boxes. During scanning, participants were shown 60 pseudo-randomized stimuli across two sessions, containing 15 object, 15 scene, 15 scenario, and 15 non-word stimuli. They were instructed to visualize the stimuli (without relying on past experiences) or count the letters for non-words. Each stimulus was presented for 10 seconds.

This was followed by a self-paced classification task, during which participants categorized their imagined content as an object, scene, scenario, or whether they were counting. On a trial-by-trial basis, they also rated their perceived vividness on a 4-point Likert scale, ranging from very vague to very vivid, via keypress. Participants responded in a self-paced manner (up to a maximum of 5 seconds).

The inter-stimulus interval ranged from 1 to 4 seconds. The experiment was run using Cogent2000 version 125 (Wellcome Centre for Human Neuroimaging, UCL, London, UK).

### Source memory task: post fMRI

After scanning, participants underwent an unexpected memory test. The stimuli presented included those previously shown during the eye-tracking and scanning sessions, as well as novel stimuli. The memory task consisted of 60 pseudo-randomized trials, with 20 trials per imagery condition (object, scene, scenario). Each condition included 10 novel and 10 old stimuli. Of the 10 old stimuli, 5 stimuli were taken from the eye-tracking, and 5 from the scanning experiment. Participants were required to indicate whether they had encountered each stimulus before and, if so, specify the context (eye-tracking or scanning) in which it was seen, or identify it as new.

### Debriefing question

We also asked participants to describe their strategies for visualizing objects, scenes, and scenarios, and whether these strategies differed between imagery types.

### Data analysis

#### Behavioural data

Behavioural data were analysed using separate repeated-measures (rm) ANOVAs, with the experimental condition (object, scene, scenario, non-word) as the within-subjects factor. For data collected during eye-tracking and scanning, we included classification accuracy, vividness ratings and reaction times as dependent variables.

For data collected after scanning, we included source (eye-tracking, scan, novel) as a second within-subjects factor and source memory accuracy (e.g., correctly recognizing a stimulus as the pre-defined category from the respective correct session) as the dependent variable.

Post-hoc comparisons were performed using paired t-tests with Bonferroni correction for multiple comparisons. Effect sizes were reported using partial eta-squared (η²p) for ANOVAs. The debriefing questions were analysed qualitatively.

Data analysis was conducted via Python libraries, including pandas, numpy, seaborn, matplotlib and pingouin.

#### Eye-tracking data analysis

For the five-second imagination duration, we analysed four eye movement parameters: (1) fixation count, (2) average fixation duration, (3) saccade count, and (4) average saccade amplitude. Data were pre-processed to remove any outliers beyond three standard deviations from the mean. For each parameter, we identified between zero and 22 individual data points (from 1140 total trials across all participants) to exclude from the analysis. No significant differences were observed between imagery categories in the outlier analysis.

Statistical analyses were conducted using rm ANOVAs for each eye movement parameter, with imagery category (object, scene, scenario) as the within-subjects factor. Post-hoc comparisons were performed using paired t-tests with Bonferroni correction for multiple comparisons. Effect sizes were reported using partial eta-squared (η²p) for ANOVAs. Data analysis was conducted via Python libraries, including pandas, numpy, seaborn, matplotlib and pingouin.

#### MRI Preprocessing

Preprocessing of MRI data was conducted using the SPM12 (Statistical Parametric Mapping 12) software package (Wellcome Trust Centre for Neuroimaging, London, UK; www.fil.ion.ucl.ac.uk/spm/) implemented in MATLAB R2019b (MathWorks, Natick, MA, USA). First, anatomical and functional images were reoriented to align with the anterior-posterior commissure axis, ensuring standardized orientation across all subjects. Field maps, comprising phase and magnitude images, were utilized to compute voxel displacement maps (VDMs). These VDMs were subsequently applied during the realignment and unwarping process to correct for geometric distortions in the echo-planar imaging (EPI) sequences. The mean functional image was co-registered to the high-resolution anatomical image. The subject’s anatomical image was segmented and spatially normalized to the T1-weighted Montreal Neurological Institute (MNI) template. Normalization parameters were then written to the functional data using the deformation fields derived from the structural image. For smoothing, a Gaussian kernel of 6 mm FWHM was applied, balancing between spatial specificity and group-level sensitivity. Rigid-body realignment was performed on the functional data to correct for head movement during scanning.

#### Partial least squares

The pre-processed fMRI data were analysed applying a partial least squares (PLS) approach, a covariance-based multivariate analysis technique with the advantage that there are no assumptions about the shape of the hemodynamic response function [35,36]. PLS employs singular value decomposition (SVD) to derive ranked latent variables (LVs) from the covariance matrix of brain activity and experimental conditions. These LVs represent patterns of brain activity that are associated with each experimental condition. Statistical significance of the LVs was assessed using permutation testing (n=500) and a value of *p* < .05 was considered significant. The reliability of each voxel contributing to the LV was assessed by bootstrapping (n=100) resulting in bootstrap ratios (BSRs).

Clusters of 10 or more voxels with a *BSR* < 2.5 (for Task-based PLS) and *BSR* < 2.0 (for Seed PLS) were considered reliable, resembling approximately *p* < .05.

Since we were interested in activity and connectivity differences between conditions, we employed a two-stage approach. In the first step, we applied a data-driven approach (mean-centred task-based PLS) to identify distinct patterns of brain activity during (1) imagery and control trials, as well as (2) in the imagery tasks only.

In the second step, we used Seed PLS analysis to assess differences in functional connectivity of the vmPFC. Seed PLS analysis investigates the relationship between the signal intensities of a predefined target region (seed voxel = vmPFC) and those of all other brain voxels, considering the influence of experimental conditions. This multivariate technique identifies patterns of brain activity that are maximally correlated with the seed region’s activity, thereby elucidating functional connectivity and its modulation by experimental manipulations. We conducted one seed PLS analysis including the three imagery tasks (mean-centred task-based Seed PLS).

#### Signal Intensity Extraction

Since episodic future simulation and episodic recall rely on similar brain regions, we extracted signal intensities from regions typically involved in autobiographical memory. The coordinates were chosen via meta-analyses maps created with Neurosynth (https://neurosynth.org/) using the term “autobiographical memory”. We included the vmPFC (MNI -4 54 -12), left/right anterior hippocampus (l: -24 -22 -18, r: 24 -14 -16), left/right posterior hippocampus (l: -28 -34 -8, r: 30 -36 -8), left/right parahippocampal gyrus (l: -24 -40 -18, r: 32 -38 -14), retrosplenial cortex (-4 -58 20), and left/right visual cortex (l: -44 -72 -2, r: 46 -72 -4). Of note, positive or negative signal intensity values extracted from PLS do not reflect fMRI activation or deactivation.

## Results

### Eye-tracking

#### Classification task

Classification accuracy did not differ between scenarios (*M* = 84%, *SD* = 37%), scenes (*M* = 75%, *SD* = 43%) and objects (*M* = 72%, *SD* = 45%, *F(*2,36) = 3.07, *p* = .06, *η²p* = .15). In cases of miss-classification, scenarios and objects were most frequently misclassified as scenes, occurring in 14% and 17% of the trials, respectively.

#### Vividness ratings

Mean vividness ratings were similar across scenarios (*M* = 1.96, SD = 0.73), objects (*M* = 2.07, *SD* = 0.79), and scenes (*M* = 2.06, *SD* = 0.75, *F*(2, 36) = 0.89, *p* = .419, *η²p* = .025). These results suggest that participants perceived similar levels of vividness across categories.

#### Reaction times

We found significant main effects of the imagery category on reaction times in the classification task (*F*(2,36) = 11.04, *p* < .001, *η²p* = .076) and the vividness rating (*F*(2,36) = 13.31, *p* < .001, *η²p* = .056). Post-hoc tests showed that scenarios were processed significantly faster than objects (Scenario classification: *M* = 1353ms; Vividness: *M* = 1385ms; Object classification: *M* = 1578ms; Vividness: *M* = 1537ms; Classification: *p_bonf_* = .024; Vividness: *p_bonf_* = .029) and scenes (Scene classification: *M* = 1717ms; Vividness: *M* = 1648ms; Classification: *p_bonf_* = .001; Vividness: *p_bonf_* < .001).

#### Eye movements

The fixation count was highest during scene construction (*M* = 3.72, *SD* = 0.97), followed by objects (*M* = 3.52, *SD* = 0.97), and scenarios (*M* = 3.50, *SD* = 0.93) with a significant main effect of imagery category (*F*(2,36) = 8.88, *p* < .001, *η²p* = .33, see **Fig.1**). Post-hoc tests showed that fixation count significantly differed between scenes and both scenarios (*p_bonf_* = .01), and objects (*p_bonf_* = .006).

**Figure 1.**
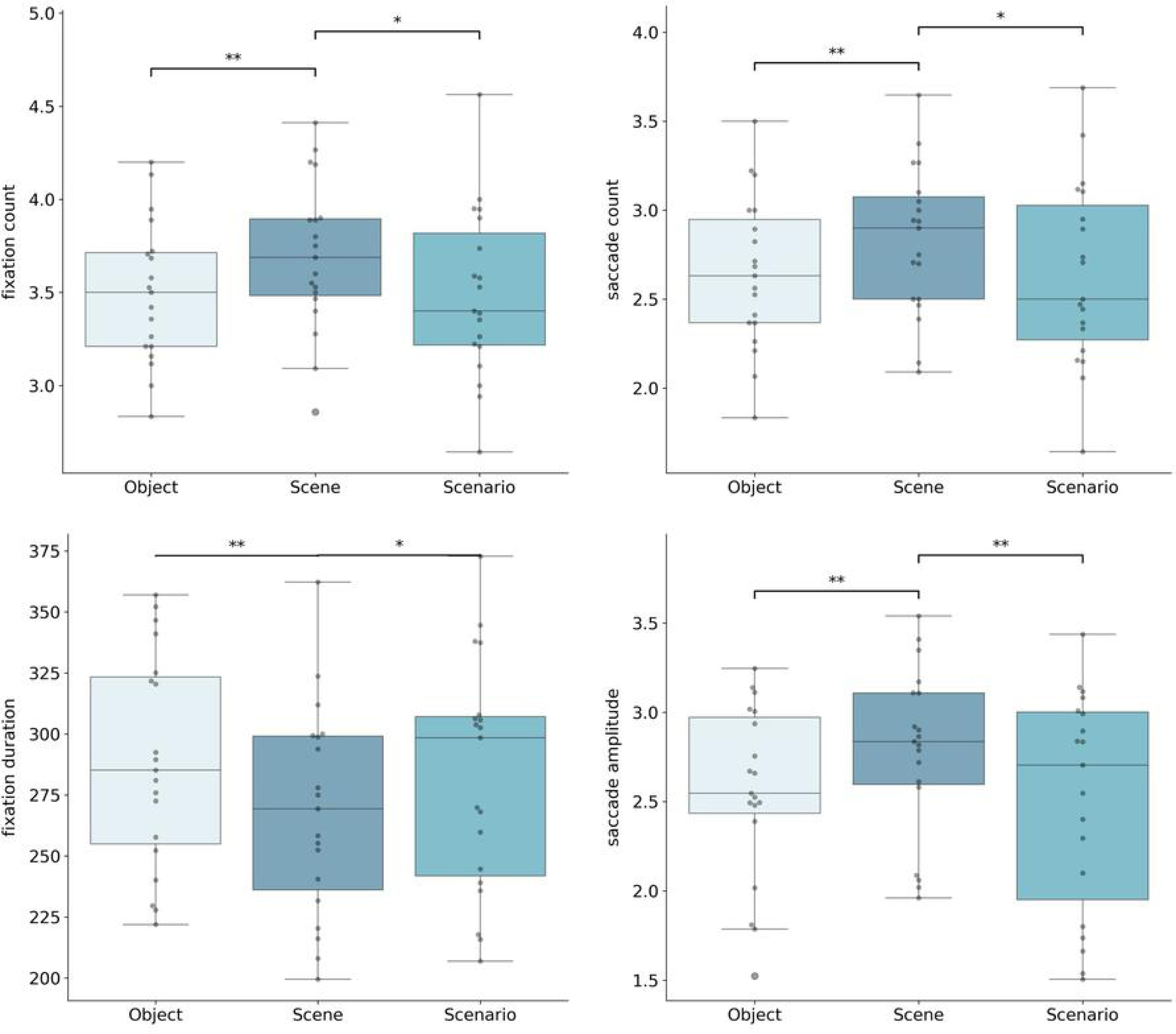
Eye movement data. Bargraphs display fixation counts / duration, saccade count / amplitude during the imagery trials *p<.05, **p<.01, ***p<.001.

The average fixation duration was longest for objects (*M* = 287.63ms, *SD* = 107.55), followed by scenarios (*M* = 282.03ms, *SD* = 102.78), and scenes (*M* = 266.93ms, *SD* = 99.44). There was a significant main effect (*F*(2,36) = 10.79, *p* < .001, *η²p* = .32), showing shorter fixation durations for scenes than both objects (*p_bonf_* = .004) and scenarios (*p_bonf_* = .01).

Scene imagery elicited more frequent saccades (*M* = 2.85, *SD* = 1.03) with larger amplitudes (*M* = 2.84°, *SD* = 1.73) compared to both scenario (saccades: *M* = 2.65, *SD* = 0.99; amplitude: *M* = 2.55°, *SD* = 1.44) and object imagery (saccades: *M* = 2.67, *SD* = 1.01; amplitude: *M* = 2.59°, *SD* = 1.50). Both saccade count (*F*(2,36) = 6.01, *p* = .006, *η²p* = .25) and saccade amplitude (*F*(2,36) = 7.15, *p* = .002, *η²p* = .28) showed significant main effects (see **Fig.1**). Post-hoc tests revealed significant differences between scene construction and both scenario and object imagery (*p_bonf_* < .05). These results replicate previous findings, showing that scene construction led to a higher number of shorter fixations, increased saccade frequency, and larger saccade amplitudes compared to object construction. [4,37,38].

### Scanning task

#### Classification task

Even though there was a significant main effect of imagery category (*F*(3,54)=3.41, *p* < .05, *n^2^p* = .16), there were no significant post-hoc tests revealing classification accuracy differences (Non-words: *M* = 91%, *SD* = 16%, objects: *M* = 90%, *SD* = 9%, scenarios: *M* = 83%, *SD* = 10%, and scenes: *M* = 79%, *SD* = 21%) (see **Fig. 2a**). Scenes were imagined as scenarios in 16% of trials, objects were imagined as either scenarios or scenes in 5%, and scenarios were imagined as scenes in 14%. Nevertheless, to account for potential misclassifications as confounding variables, we repeated our fMRI analyses based on the individual subject response categories.

**Figure 2.**
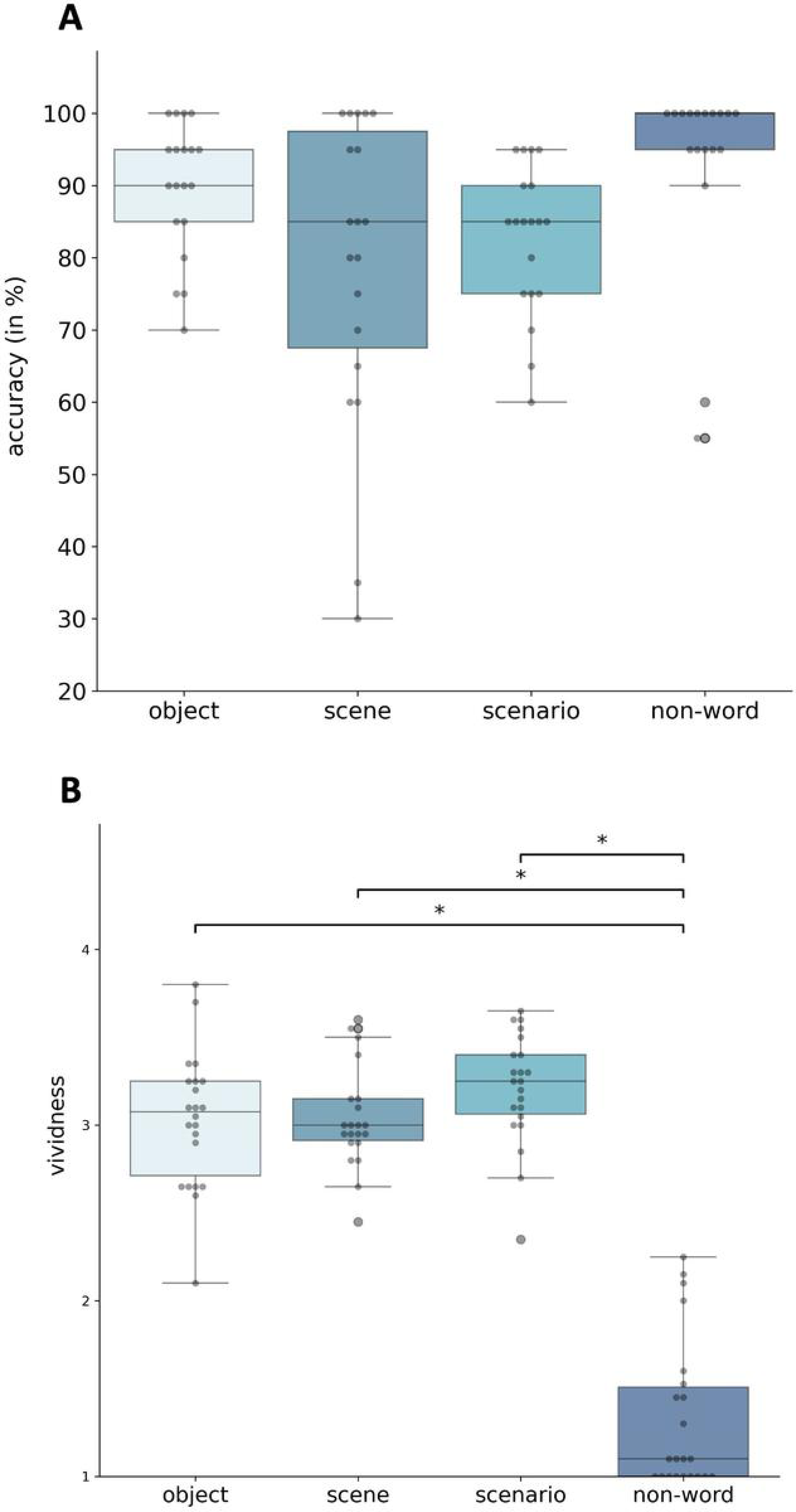
Scanning task. A: Classification accuracy (in percent) B: Vividness ratings. Higher scores resemble higher vividness ratings. *p<.05, **p<.01, ***p<.001

#### Vividness rating

Vividness ratings (see **Fig.2b**) were expectedly lowest for non-words (*M* = 1.31, *SD* = 0.54), and equally high for objects (*M* = 2.98, *SD* = 0.71), scenes (*M* = 3.03, *SD* = 0.69) and scenarios (*M* = 3.18, *SD* = 0.74) with a significant main effect of category (*F*(3,54) = 143.62, *p* < .001, *n^2^p* = .89). Post-hoc analyses confirmed the expected significant differences between non-words and the imagery conditions (*p’s* < .05). Since vividness did not differ between the imagery categories, the rating was not considered as a mediating variable in the analysis of brain activity data in the multivariate analysis.

#### Reaction times

Mean reaction times for classification of the stimuli was fastest for non-words (M = 924.05ms, SD = 493.12), followed by objects (M = 1175.22ms, SD = 614.72) scenarios (M = 1228.64ms, SD = 659.74), and scenes (M = 1467.44, SD = 794.93) with a significant main effect of category (*F*(3,54) = 17.09, *p* < .001, *n^2^p* = .49). Post-hoc tests revealed significant differences between non-words and imagery stimuli (*p_bonf_’s* < .05), as well as between scenes and both objects (*p_bonf_* = .004) and scenarios (*p_bonf_* = .001).

Mean reaction times for vividness showed only mild variation across categories: non-words (M = 817.35ms, SD = 539.59), scenarios (M = 1105.71ms, SD = 726.25), scenes (M = 1112.12ms, SD = 705.50), and objects (M = 1160.85ms, SD = 732.54). While a significant main effect was observed (*F*(3,54) = 7.4, *p* = .002, *n^2^p* = .29), all post-hoc comparisons failed to reach statistical significance (*p_bonf_* > .05). This suggests that, despite overall differences in reaction times, the categories did not vary systematically on a behavioural level.

#### Post-scan source memory task

Source memory is defined as recognition accuracy, i.e., correct classification of a debriefing stimulus into the pre-defined imagery category (a scene stimulus as scene) and identification of correct session (a stimulus from the scanning session as from the scanning session). This varied across imagery categories and sessions (see **Fig. 3**).

**Figure 3.**
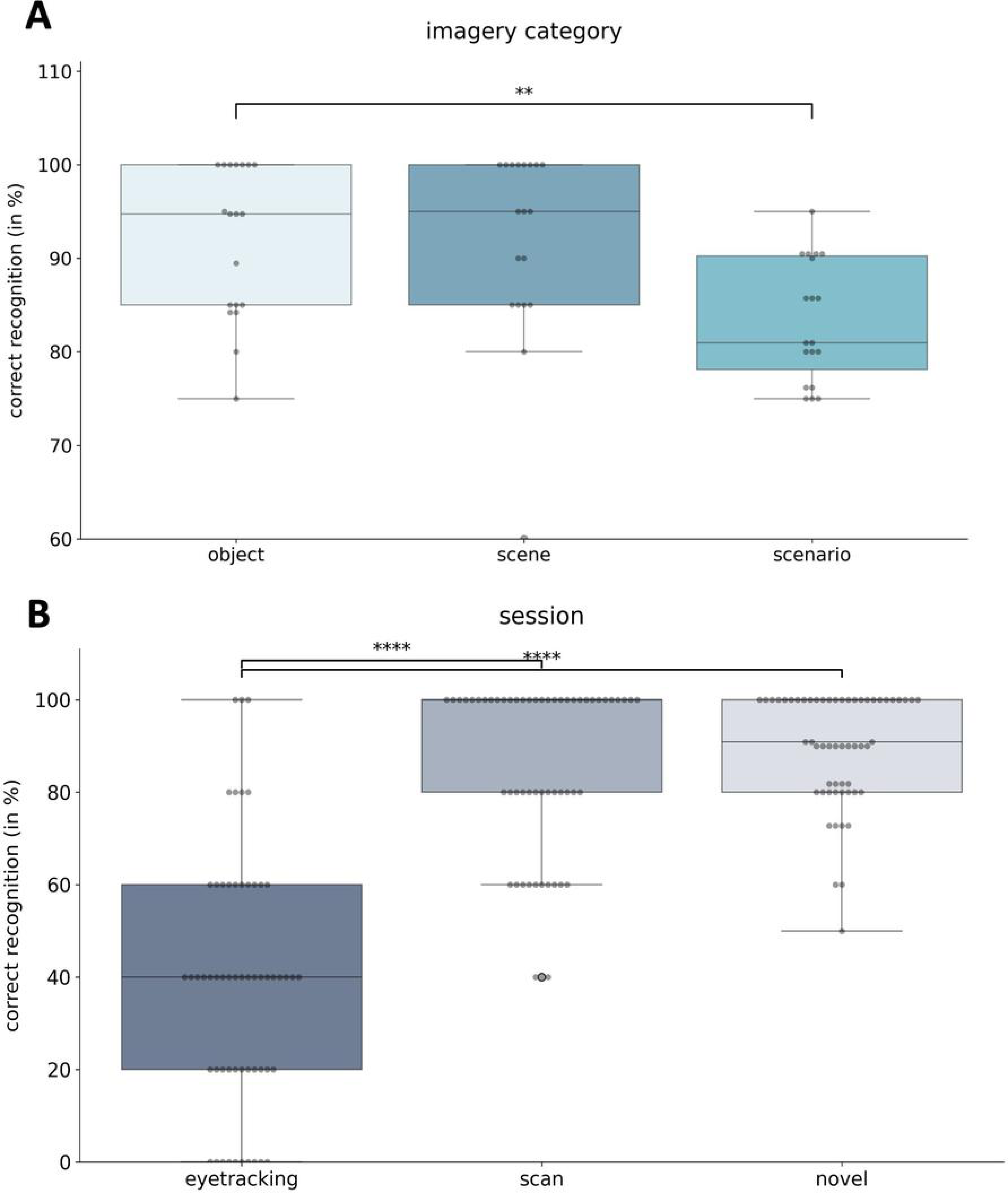
Post-scan Source Memory Task. A: Bar graph showing the correct recognition (in percent) of stimuli as object, scene or scenario stimuli. B: Bar graphs showing the correct recognition (in percent) of stimuli from their respective session. *p<.05, **p<.01, ***p<.001

#### Imagery category

Scenarios were correctly recognized in 83% (SD = 6%), scenes in 92% (SD = 10%), and objects in 92% (SD = 8%) of all trials, *F*(2,36) = 6.70, *p* = .005, *n^2^p* = .35. Post-hoc tests revealed significant differences between scenarios and objects (*p_bonf_* = .003).

#### Session

Recognition accuracy was lowest for stimuli from the eye-tracking session (M = 38%, SD = 27%), followed by stimuli from the scanning session (M = 85%, SD = 19%), and highest for novel stimuli (M = 90%, SD = 10%), *F*(2, 36) = 60.02, *p* = < .001, *η^2^p* = .77). Post-hoc tests revealed significant differences between eye-tracking and both the scanning session and novel stimuli (*p_bonf_’s* < .001). There was no significant interaction effect (p = .15) Importantly, when analysing recognition accuracy only for the stimuli from the scanning session, no significant differences were found (*p* = .89). We conclude that encoding/memory load during scanning did not differ between the categories and thus cannot account for the differences in the fMRI findings.

#### Debriefing question

After scanning, all participants described significant differences between imagining objects, scenes and scenarios. Most participants reported that imagining objects began with a basic shape, which was then enriched with details - some perceived as 2D and others as 3D. Objects were typically imagined against a white background with some mentioning that the object slowly rotated in their mind’s eye. In contrast, scenes were initially constructed as a 3D spatial framework, with individual elements and details integrated progressively to form a coherent, static image, resembling a photograph. A scenario, however, was described as a temporally extended, dynamically unfolding sequence, characterized by movement of both people and oneself. Importantly, scenarios were always embedded within a spatially coherent context, which evolved over time, reinforcing the movie-like nature of these mental events. These descriptions support our view, that imagining objects, scenes or scenarios resemble qualitatively different cognitive experiences. Only two participants struggled with generating new content and reported relying on their memories in some of the trials.

### Data-driven PLS

#### Mean-centred PLS with all four conditions

We used mean-centred task-based PLS analysis to assess neural differences between the four conditions. This analysis showed significant differences in brain patterns associated with imagery and the control condition (*p* < .001). Positive brain scores (or saliences) were associated with regions in which the BOLD signal was greater for imagery. Negative brain scores (or saliences) were associated with regions in which the BOLD signal was greater for the control condition.

Within the imagery pattern, scenario construction had a stronger influence than scene and object construction, which made comparable contributions. This pattern encompassed bilateral vmPFC, anterior and posterior hippocampus, parahippocampal and lingual gyri, right precuneus, and bilateral middle occipital gyri. The control condition was associated with increased BOLD signal across a distributed fronto-parieto-occipital and lateral temporal brain pattern (see **Fig. 4**, supplementary tables for cluster reports).

**Figure 4.**
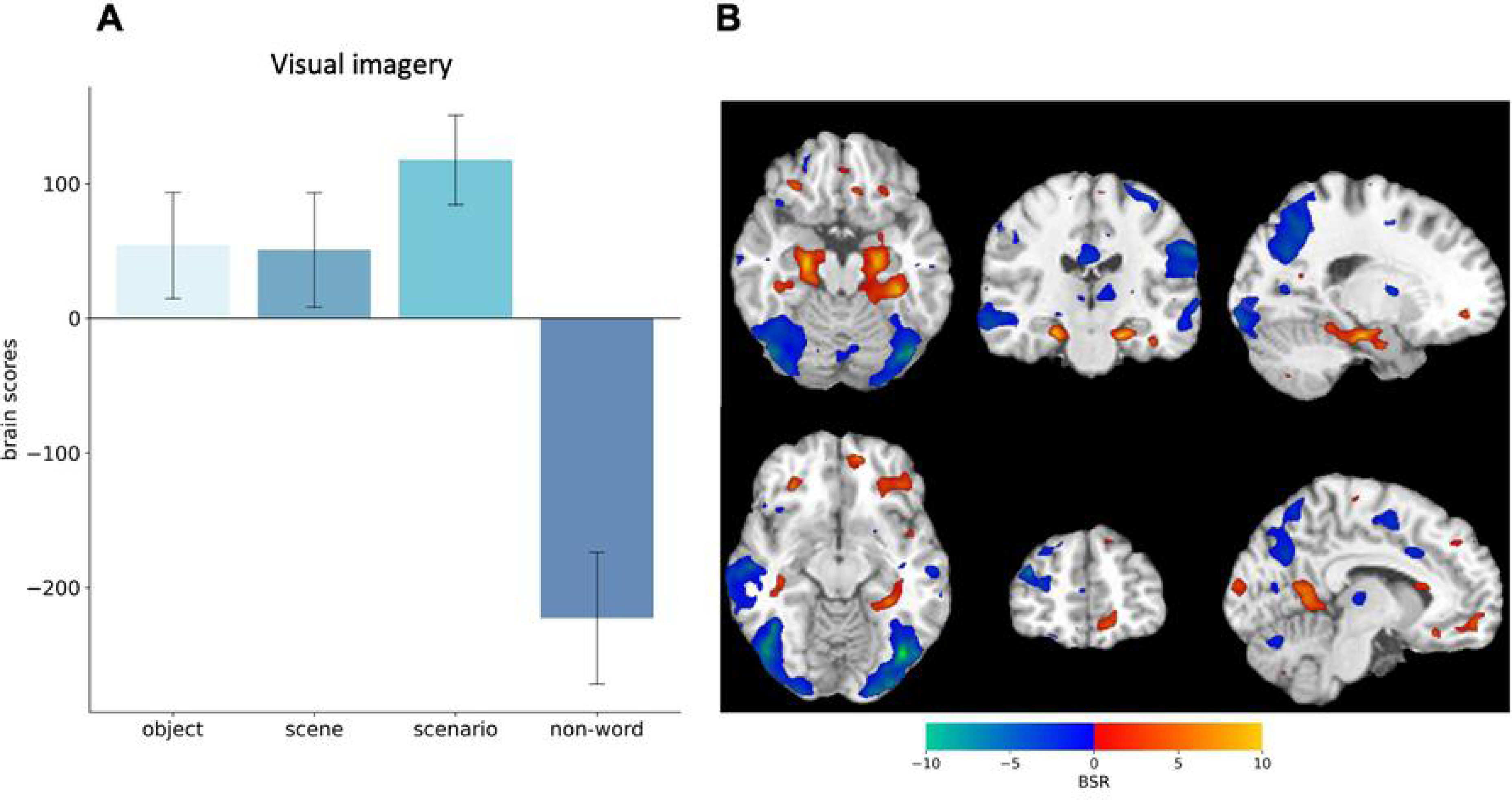
Task-based PLS results from four conditions. (A) Brain scores associated with visual imagery (LV1) differentiate object, scene, and scenario construction from the control task. Bar graphs display means with 95% bootstrapped confidence intervals. (B) Bootstrap Ratios (BSR) are displayed on a single-subject 1T template ni standard space. Warm colors indicate increased activity during imagery tasks, while cool colors represent increased activity during the control condition. The statistical map is thresholded at BSR = ±2.5.

**Figure 5.**
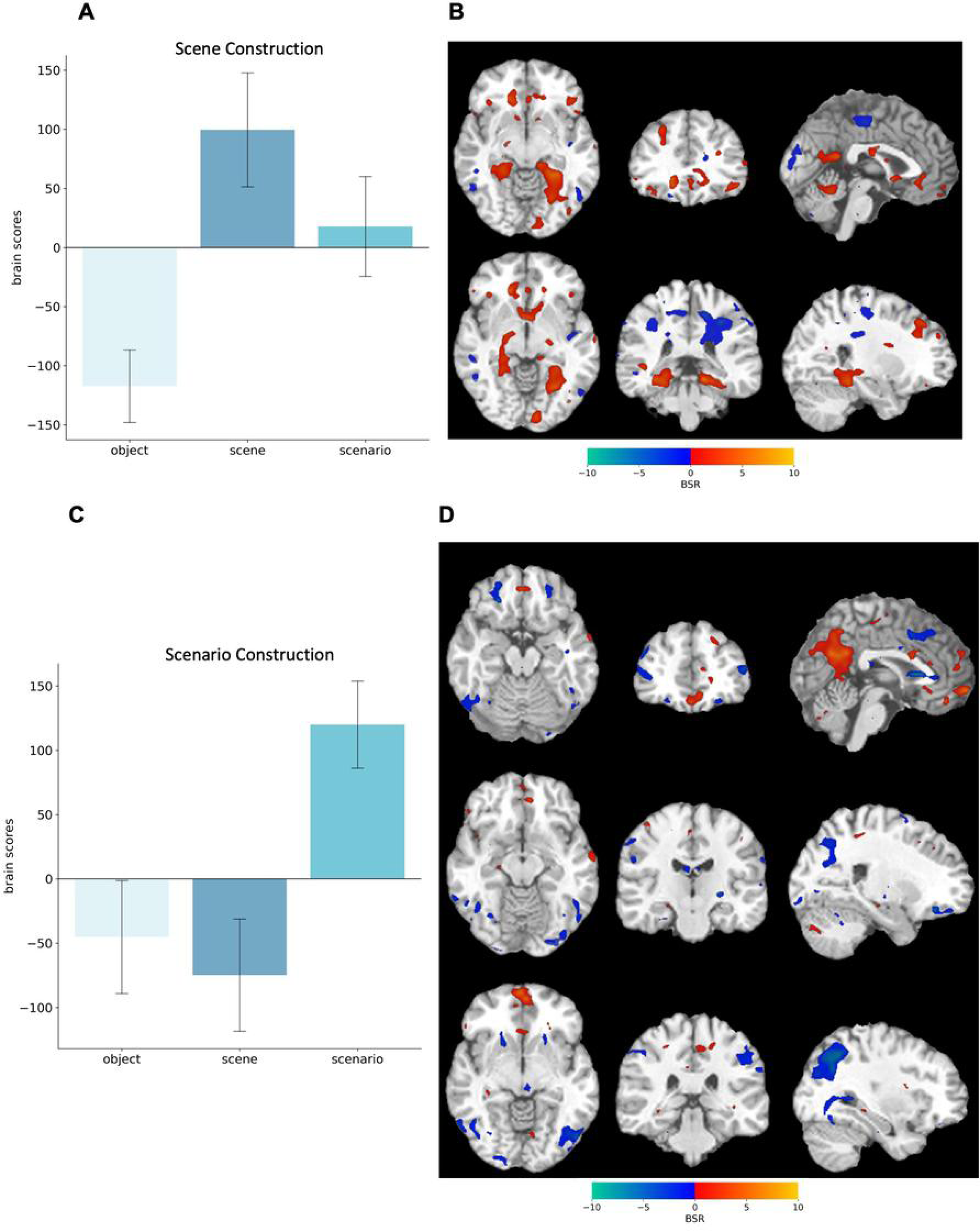
Task-based PLS results from the imagery conditions. (A) Brain scores associated with the first latent variable (LV1) differentiating scene from object construction, and (C) the second LV2 differentiating scenario from both object and scene construction. Bar graphs display means with 95% boot-strapped confidence intervals. (B) Bootstrap Ratios (BSR) are displayed on a single-subject 1T template ni standard space. Warm colours reflect activity during scene construction, whereas cool colours reflect activity during object construction. (D) Warm colours reflect activity during scenario construction, whereas cool colours reflect activity during object and scene construction. The statistical map is thresholded at BSR = ±2.5.

#### Mean-centred PLS with all three imagery conditions

Next, we repeated the task-based PLS analysis without the control condition to focus on differences between the imagery conditions. We found two significant LVs, representing (1) Scene construction and (2) Scenario construction.

### Scene Construction

The first LV (*p* = .010) differentiated scene (positive brains scores) from object construction (negative brain scores). Scenario construction did not contribute to this pattern (confidence intervals including zero, see **Fig.5a**). During scene construction, there was increased bilateral activity in the vmPFC, parahippocampal gyrus, lingual gyrus, and left lateral occipital lobe (see **Fig.5b**). In contrast, object construction was associated with fronto-parietal brain areas, cuneus, and posterior cingulate cortex.

### Scenario Construction

The second LV (*p* = .026) differentiated scenario (positive brain scores) from scene and object construction (negative brain scores, see **Fig.5c**). Scenario construction was supported by strong bilateral activation in the vmPFC, as well as in the left dorsolateral frontal areas, right anterior and posterior hippocampus, and bilateral posterior cingulate cortex, anterior precuneus, and angular gyrus. In contrast, scene / object construction was associated with bilateral dorsal and lateral frontal cortex, bilateral posterior precuneus, bilateral fusiform gyri, right lingual gyrus, and inferior occipital lobe (see **Fig.5d**).

#### Regional signal intensities

We extracted signal intensities from ten a priori selected regions of interest (see **Fig. 6**) to examine activity patterns across imagery conditions. Repeated-measures ANOVAs revealed condition-specific engagement patterns across several regions. In agreement with our primary hypothesis, the vmPFC showed significant modulation by condition (*F*(2, 36) = 10.76, *p* < .001, *n^2^p* = .37), with stronger engagement during scenario construction compared to both scene (*p_holm_* = .006) and object construction (*p_holm_* = .002).

**Figure 6.**
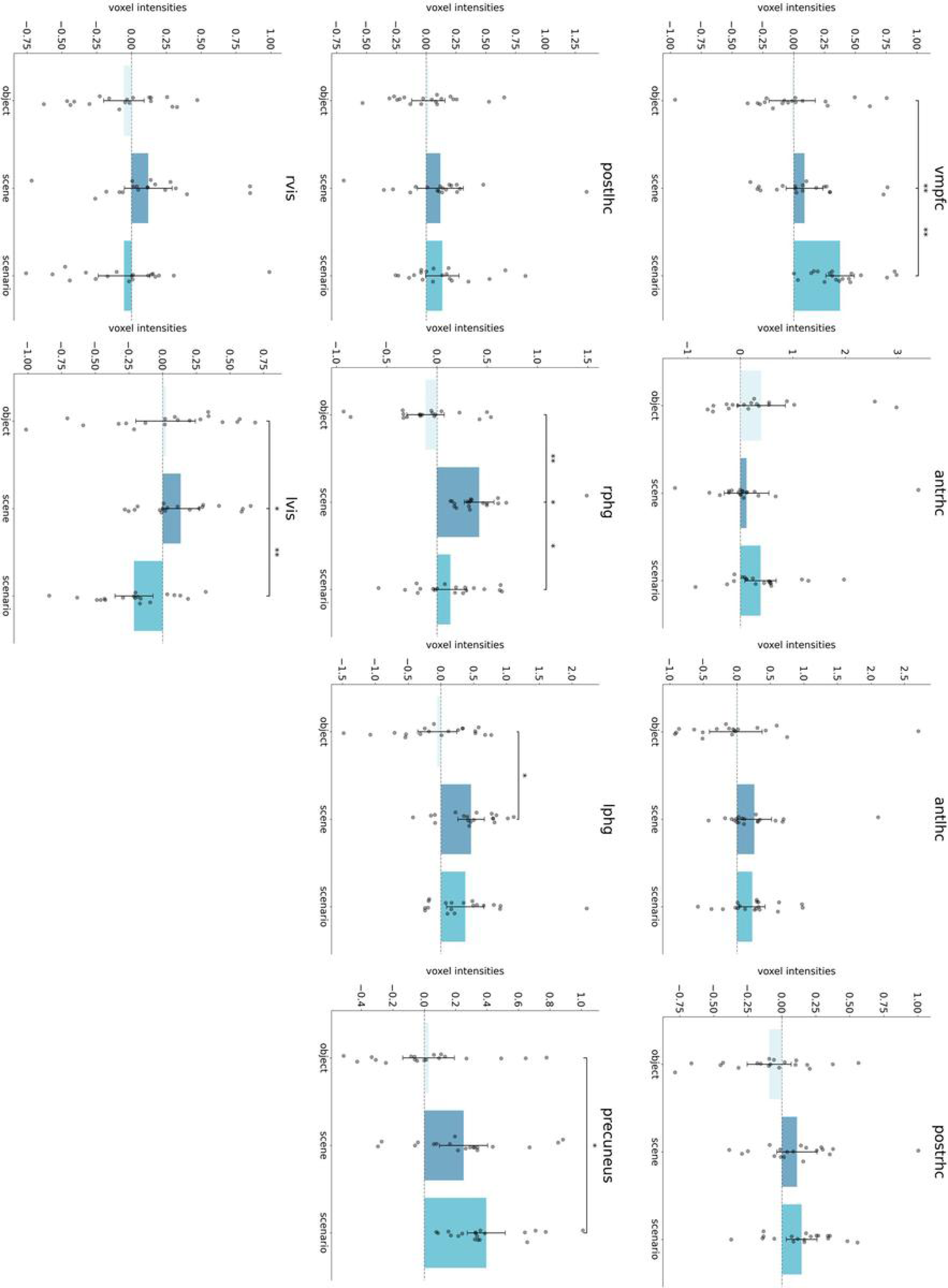
Extracted signal intensities from brain regions associated with autobiographical memory taken from NeuroSynth. Violin plots display the distribution of individual signal intensities. Box plots display the median and the interquartile range of voxel intensities. *p<.05, **p<.01, ***p<.001

Medial temporal lobe regions also showed marginally significant to significant condition-specific effects. The right posterior hippocampus (*F*(2, 36) = 4.29, *p* = .021, *η²p* = .19) and right parahippocampal gyrus (*F*(2, 36) = 9.53, *p* < .001, *η²p* = .35) both demonstrated stronger engagement during scenario than object construction (*p_holm_* = .051 and *p_holm_* = .046, respectively). Additionally, both regions showed greater activation during scene than object construction (*p_holm_* = .051 and *p_holm_* = .003, respectively). The left parahippocampal gyrus exhibited a different pattern (*F*(2, 36) = 4.67, *p* = .01, *η²p* = .21), with scenes eliciting greater activation than both scenarios (*p_holm_* = .052) and objects (*p_holm_* = .026). We found no significant differences between conditions in the anterior hippocampi or in the posterior segment of the left hippocampus.

Within the precuneus, we observed a gradation pattern (*F*(2, 36) = 5.62, *p* = .007, *η²p* = .24), in which scenario stimuli elicited stronger engagement than object stimuli (*p_holm_* = .01). In the visual perceptual cortex, hemispheric differences emerged. While no significant condition effects were observed in the right hemisphere, the left hemisphere showed condition-dependent activation (*F*(2, 36) = 7.95, *p* = .001, *η²p* = .30), with stronger engagement during object (*p_holm_* = .019) and scene (*p_holm_* = .001) construction compared to scenario construction.

#### Functional vmPFC-neocortical connectivity during visual imagery

Since our main goal was to assess vmPFC contributions to imagery construction, we conducted a seed PLS analyses with the peak vmPFC voxel (MNI: -4 54 -12) being the seed. Two significant LV’s (see **Fig. 7** and supplementary Tables 2a,b) were identified, which showed a gradient in how the vmPFC functionally connects with the brain network.

**Figure 7.**
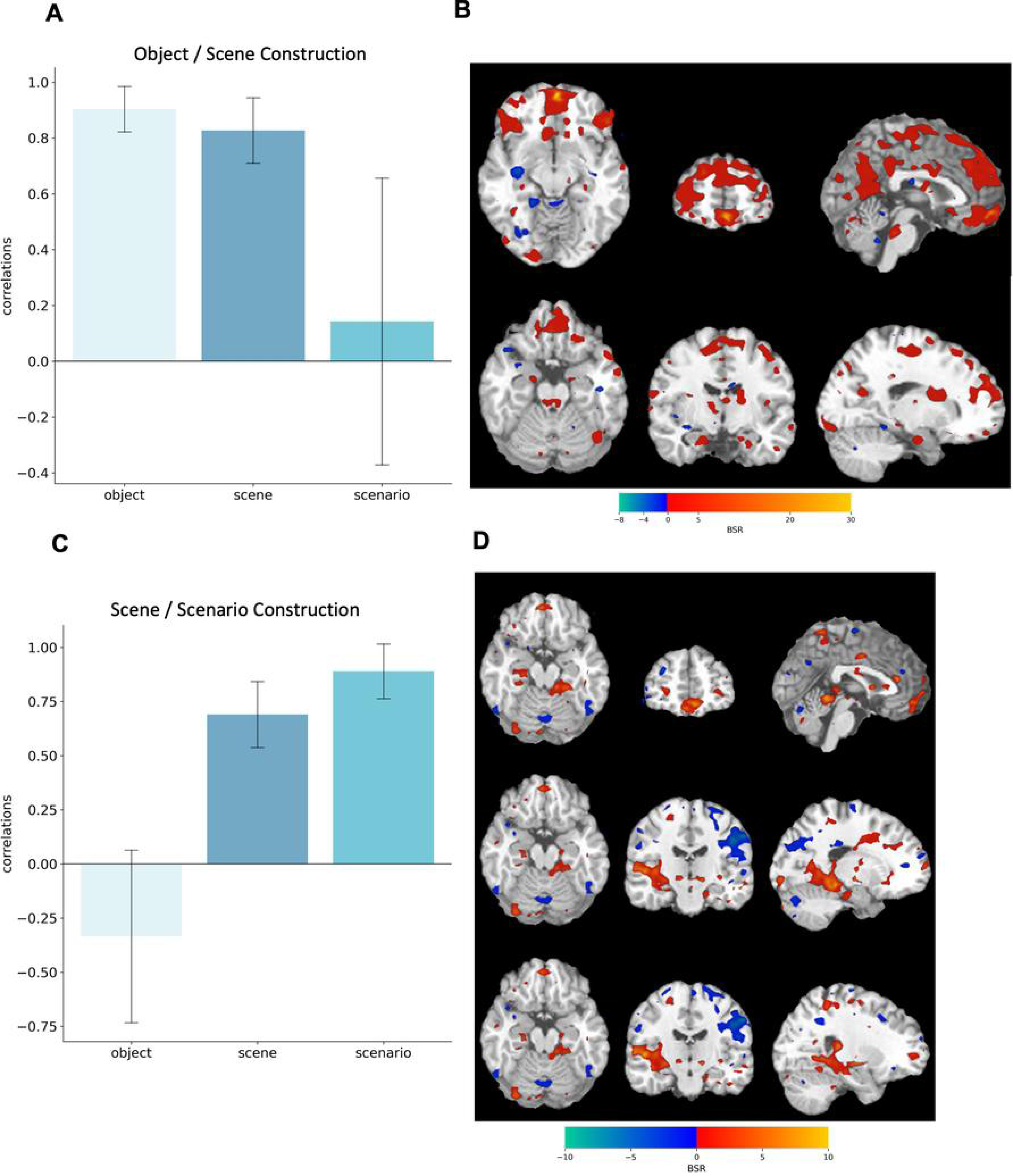
Seed-PLS results. Correlations associated with the significant latent variables LV1 (A) & LV2 (C) differentiating object, scene and scenario imagery. Bar graphs display means with 95% boot-strapped confidence intervals. (C) Bootstrap Ratios (BSR) are displayed on a single-subject T1 template in standard space. Warm colours reflect activity during scene and object construction, whereas cool colours reflect activity during object construction. (B) Warm colours reflect connectivity during object and scene construction. (D) Warm colors reflect connectivity during scene and scenario construction. The statistical map is thresholded at BSR = ±2.0.

The first LV (*p* < .001) showed a strong negative connectivity during object (*r* = -0.90) and scene imagery (*r* = -.83), but not during scenario imagery (*r* = -.14, see **Fig. 7a**). Notably, signal intensity changes in the vmPFC were correlated with changes in bilateral fronto-parietal networks, anterior medial and posterior hippocampus, as well as precuneus and retrosplenial cortex (see **Fig. 7b**).

The second LV (*p* = .030) differentiated scene and scenario construction from object construction (see **Fig. 7c**). The strongest effect is observed in scenarios (*r* = .89), followed by scenes (*r* = .69), with objects showing an inverse relation (*r* = -.33). Regions exhibiting reliable positive functional connectivity during scenario and scene construction included bilateral anterior and posterior hippocampi, right parahippocampal cortex, and bilateral fusiform and lingual gyri. In contrast, object construction was associated with stronger functional connectivity with frontotemporal and frontoparietal networks (see **Fig. 7d**).

## Discussion

This study utilized high-resolution 7T MRI to investigate the neural substrates supporting three forms of mental imagery—object, scene, and scenario construction. We proposed a hierarchical organization in which the posterior neocortex provides visual-perceptual details, the hippocampus supports spatial scene construction, and the ventromedial prefrontal cortex (vmPFC) integrates these components into temporally and spatially structured scenarios.

This study directly compares three forms of mental imagery: single objects without a spatial background, static isolated scenes, and naturalistically evolving mental scenarios, while previous research has predominantly focused on the neural underpinnings of scene and object imagery in isolation, leaving a critical gap in understanding the specific features that drive neural differences between these forms of mental imagery [26,39–44]. We hypothesized that posterior cortices would support all imagery types by providing detailed visual information about isolated objects and scenes comprising scenarios. The hippocampus was expected to contribute to scene and scenario construction by creating the spatial context for scenes and scenarios, while the vmPFC was hypothesized to be most engaged in temporally extended scenarios, serving as an integrative hub.

### Common neural pattern for imagery-rich mental events

The activation pattern distinguishing imagery conditions from the low-level control task (letter counting in non-words) included the vmPFC, anterior / posterior hippocampus, and the visual-perceptual cortex – a network consistently implicated in mental imagery across numerous studies [23–25,45–49].

### vmPFC: Temporal Integration and Scenario Construction

Our findings strongly support the hypothesis that the vmPFC plays a hierarchical role in visual imagery, with increasing engagement as imagery becomes more complex and contextually rich (objects → scenes → scenarios). This was demonstrated by multiple analyses: (1) In the mean-centred PLS (including all conditions), scenario construction showed the strongest association to a neural pattern that included the vmPFC, anterior / posterior hippocampus, and visual-perceptual cortices - regions consistently linked to scene processing. (2) In the mean-centred PLS (imagery conditions only), we identified a significant latent variable that distinguished scenario from scene construction, showing that scenario construction was associated with a distinct neural pattern that prominently involved the vmPFC. (3) A priori selected ROI analyses revealed that the vmPFC was more strongly engaged during scenario than both scene and object construction. (4) Functional connectivity analysis with the vmPFC as a seed revealed that during scenario and scene construction, the vmPFC exhibited stronger positive connectivity with a widespread brain network, including the hippocampus and visual-perceptual cortices, compared to object construction. The distinct connectivity pattern during scenario construction suggests the vmPFC may serve as an integrative hub, potentially reflecting its role in integrating spatial, contextual, and possibly autobiographical or semantic information into coherent mental representations.

In line with our findings, previous research has highlighted the vmPFC’s role in constructing temporally extended mental simulations. First, patients with vmPFC lesions show impairments in autobiographical memory retrieval and future thinking, which require integrating multiple scenes into coherent, unfolding scenarios, while single scene processing remains intact [7,8,15,50]. Furthermore, recent MEG studies have provided evidence that the vmPFC exerts a driving influence on the hippocampus during tasks that require the temporal unfolding of mental events [19–21].

Our results extend these findings by directly comparing scene and scenario construction within the same study, revealing that the vmPFC is significantly more engaged when constructing dynamic, temporally structured scenarios than static scenes.

### Hippocampus: Scene Construction

Our findings indicate that the hippocampus and surrounding medial temporal lobe structures support scenario and scene construction to a greater extent than object construction. The mean-centred PLS revealed two significant LVs: LV1 showed greater hippocampal/parahippocampal activation for scenes vs. objects, while LV2 showed greater hippocampal activation for scenarios vs. objects. ROI analyses confirmed this pattern, particularly in the right parahippocampal gyrus and the left parahippocampal gyrus.

Previous research connects scene construction to the hippocampus [3–5,7,12,27], its anterior segment [23–26]. However, the pattern was not corroborated by the ROI analysis, which failed to reveal a strong preference of the anterior hippocampus for scene versus object construction. This discrepancy may be attributed to our approach of preselecting ROIs based on the Neurosynth database, which potentially did not capture the peak of difference in our sample. It is worth noting that a subsequent analysis of this dataset, which extends beyond the scope of the current investigation, will examine hippocampal subfields in greater detail to elucidate their specific contributions to the construction of mental imagery-rich events.

### Posterior neocortex: Sensory-Perceptual Support

The posterior neocortex was engaged in all imagery conditions, as shown in the first PLS analysis contrasting all imagery conditions to the low-level baseline. The areas comprised the retrosplenial cortex, precuneus, lingual, and fusiform gyrus, as well as middle occipital gyrus. We hypothesised previously that the posterior neocortex provides visual-perceptual elements to any type of mental imagery-rich events, such as colour, shape and movement details [51–56]. Since object construction necessitates focussed attention on such details, it is fitting that the ROI analysis of the left visual cortex revealed greater activation during object construction relative to other types of mental imagery.

Higher engagement of the precuneus during scenario construction, as indicated by the PLS and ROI analyses is likely due to its role in self-referential processing, temporal and spatial integration of complex information, which is essential for mentally constructing spatiotemporally coherent mental simulations [57,58].

### Behavioural and Eye Movement Differences in Response to Cognitive Demands During Visual Imagery

Lastly, the behavioural and oculomotor data provide further critical insights into the constituents of scenario construction distinguishing it from scene and object construction. The imagery conditions were carefully designed and piloted to ensure that vividness, construction demands, and word length were consistent across conditions, isolating the precise mental content as the key difference.

During object construction, participants reported focusing on a single, detailed object devoid of spatial context, confirmed by less frequent smaller-amplitude saccades and long fixations compared to scenes. In contrast, scene construction was described to involve immediately generating a spatial layout, subsequently filled with multiple visuospatial details, reflected in frequent, short fixations, and large-amplitude saccades - indicative of constructing a more spatially extended mental space. This pattern aligns with prior findings on scene construction [4,37,38,59].

While temporally extended scenarios were remembered and visualized as vividly and accurately as scenes and objects, participants described scenario construction as a sequential and combinatory process, starting with rapid spatial layout generation (like scenes) and followed by the construction of individual foreground objects integrated into an unfolding narrative. Interestingly, despite the similarity in neuronal activation patterns between scenarios and scenes, eye-tracking data revealed that scenarios and objects triggered more similar fixation durations and saccade amplitudes than scenes. This apparent discrepancy may be explained by the dual processing demands of scenario construction: object-focused visual attention, as captured by eye-tracking metrics [59–61], and scene-based integration, as reflected in fMRI data. Together, these findings highlight the complex interplay between visual attention and neural mechanisms in constructing temporally extended mental events, underscoring the unique cognitive demands of scenario construction.

## Conclusion

Our study provides a comprehensive neural framework for how the brain constructs naturalistic, imagery-rich events. While the posterior neocortex provides the perceptual details necessary for constructing single objects, scenes, and scenarios, the hippocampus supports the spatial layout essential for scene and scenario construction. The vmPFC, in turn, plays a progressively integrative role, with its greatest engagement occurring during scenario construction, where it combines and elaborates relevant information to form a coherent mental representation. These findings refine current models of mental imagery and offer a foundation for investigating its impairments in clinical conditions further.

## Acknowledgements

The authors would like to thank A. Ruehling for excellent technical assistance during the acquisition of the imaging data. The imaging experiments were performed at the German Center for Neurodegenerative Diseases (DZNE), Bonn, Germany.

## Data availability statement

The data supporting this study are available from the corresponding author upon reasonable request and following approval of an appropriate ethics proposal.

## Funding

This research was supported by the Hertie Network of Excellence in Clinical Neuroscience. Work in C.M.’s lab is further financed by internal research funding of the Faculty of Medicine (BONFOR), University Hospital Bonn, by the Federal Ministry of Education and Research (BMBF) within the framework of the funding programme ACCENT (funding code 01EO2107) and by the Deutsche Forschungsgemeinschaft (DFG, German Research Foundation, MC 244/3-1). J.T. received an Argelander Mobility Grant from the University Bonn.

## Supporting Information

**Table1a**. Cluster report for the first latent variable from the mean-centered Task-based PLS with the imagery conditions

**Table 1b.** Cluster report for the second latent variable from the mean-centered Task-based PLS with the imagery conditions

**Table 2a.** Cluster report from the first latent variable from the mean centered task-based Seed PLS

**Table 2b.** Cluster report from the second latent variable from the mean centered task-based Seed PLS

## Notes

### Competing Interest Statement

The authors have declared no competing interest.

## References

[1] Kim S, Dede AJO, Hopkins RO, Squire LR. Memory, scene construction, and the human hippocampus. Proc Natl Acad Sci U S A 2015;112:4767–72. 10.1073/PNAS.1503863112/SUPPL_FILE/PNAS.201503863SI.PDF.

[2] Hassabis D, Kumaran D, Vann SD, Maguire EA. Patients with hippocampal amnesia cannot imagine new experiences. Proc Natl Acad Sci U S A 2007;104:1726–31. 10.1073/pnas.0610561104.

[3] Maguire EA, Mullally SL. The hippocampus: A manifesto for change. J Exp Psychol Gen 2013;142:1180–9. 10.1037/a0033650.

[4] McCormick C, Maguire EA. The distinct and overlapping brain networks supporting semantic and spatial constructive scene processing. Neuropsychologia 2021;158. 10.1016/J.NEUROPSYCHOLOGIA.2021.107912.

[5] McCormick C, Dalton MA, Zeidman P, Maguire EA. Characterising the hippocampal response to perception, construction and complexity. Cortex 2021;137:1–17. 10.1016/J.CORTEX.2020.12.018.

[6] McCormick C, Rosenthal CR, Miller TD, Maguire EA. Deciding what is possible and impossible following hippocampal damage in humans. Hippocampus 2017;27:303–14. 10.1002/HIPO.22694.

[7] McCormick C, Ciaramelli E, De Luca F, Maguire EA. Comparing and Contrasting the Cognitive Effects of Hippocampal and Ventromedial Prefrontal Cortex Damage: A Review of Human Lesion Studies. Neuroscience 2018;374:295–318. 10.1016/j.neuroscience.2017.07.066.

[8] Bertossi E, Ciaramelli E. Ventromedial prefrontal damage reduces mind-wandering and biases its temporal focus. Soc Cogn Affect Neurosci 2016;11:1783–91. 10.1093/SCAN/NSW099.

[9] McCormick C, Rosenthal CR, Miller TD, Maguire EA. Mind-Wandering in People with Hippocampal Damage. The Journal of Neuroscience 2018;38:2745. 10.1523/JNEUROSCI.1812-17.2018.

[10] Luelsberg F, Krakau S, Chaieb L, Witt JA, von Wrede R, Fell J, et al. Neuropsychological features of mind wandering in left-, right- and extra temporal lobe epilepsy. Seizure 2022;95:50–5. 10.1016/J.SEIZURE.2021.12.013.

[11] Costi S, Stone J, Fell J. Is the Hippocampus a Potential Target for the Modulation of Mind Wandering in Major Depression? Front Psychiatry 2018;9:363. 10.3389/FPSYT.2018.00363.

[12] Hassabis D, Kumaran D, Vann S, Maguire E. Patients with hippocampal amnesia cannot imagine new experiences. Proceedings of the National Academy of Sciences 2007. 10.1073/PNAS.0610561104.

[13] Rosenbaum RS, Moscovitch M, Foster JK, Schnyer DM, Gao F, Kovacevic N, et al. Patterns of autobiographical memory loss in medial-temporal lobe amnesic patients. J Cogn Neurosci 2008;20:1490–506. 10.1162/JOCN.2008.20105.

[14] Rayner G, Antoniou M, Jackson G, Tailby C. Compromised future thinking: another cognitive cost of temporal lobe epilepsy. Brain Commun 2022. 10.1093/BRAINCOMMS/FCAC062.

[15] Kurczek J, Wechsler E, Ahuja S, Jensen U, Cohen NJ, Tranel D, et al. Differential contributions of hippocampus and medial prefrontal cortex to self-projection and self-referential processing. Neuropsychologia 2015;73:116. 10.1016/J.NEUROPSYCHOLOGIA.2015.05.002.

[16] Karapanagiotidis T, Bernhardt BC, Jefferies E, Smallwood J. Tracking thoughts: Exploring the neural architecture of mental time travel during mind-wandering. Neuroimage 2017;147:272–81. 10.1016/J.NEUROIMAGE.2016.12.031.

[17] Lieberman MD, Straccia MA, Meyer ML, Du M, Tan KM. Social, self, (situational), and affective processes in medial prefrontal cortex (MPFC): Causal, multivariate, and reverse inference evidence. Neurosci Biobehav Rev 2019;99:311–28. 10.1016/J.NEUBIOREV.2018.12.021.

[18] Ciaramelli E, De Luca F, Monk AM, Mccormick C, Maguire EA. What “wins” in VMPFC: Scenes, situations, or schema? 2019. 10.1016/j.neubiorev.2019.03.001.

[19] McCormick C, Barry DN, Jafarian A, Barnes GR, Maguire EA. vmPFC Drives Hippocampal Processing during Autobiographical Memory Recall Regardless of Remoteness. Cerebral Cortex (New York, NY) 2020;30:5972. 10.1093/CERCOR/BHAA172.

[20] Monk AM, Barnes GR, Maguire EA. The Effect of Object Type on Building Scene Imagery—an MEG Study. Front Hum Neurosci 2020;14:592175. 10.3389/FNHUM.2020.592175/BIBTEX.

[21] Monk AM, Dalton MA, Barnes GR, Maguire EA. The role of hippocampal– ventromedial prefrontal cortex neural dynamics in building mental representations. J Cogn Neurosci 2020;33:89–103. 10.1162/JOCN_A_01634.

[22] Monk AM, Barry DN, Litvak V, Barnes GR, Maguire EA. Watching Movies Unfold, a Frame-by-Frame Analysis of the Associated Neural Dynamics. ENeuro 2021;8. 10.1523/ENEURO.0099-21.2021.

[23] Angeli PA, DiNicola LM, Saadon-Grosman N, Eldaief MC, Buckner RL. Specialization of the human hippocampal long axis revisited. Proc Natl Acad Sci U S A 2025;122. 10.1073/PNAS.2422083122.

[24] Zeidman P, Maguire EA. Anterior hippocampus: the anatomy of perception, imagination and episodic memory. Nat Rev Neurosci 2016. 10.1038/NRN.2015.24.

[25] Zeidman P, Lutti A, Maguire EA. Investigating the functions of subregions within anterior hippocampus. Cortex 2015;73:240. 10.1016/J.CORTEX.2015.09.002.

[26] Leelaarporn P, Dalton MA, Stirnberg R, Stöcker T, Spottke A, Schneider A, et al. Hippocampal subfields and their neocortical interactions during autobiographical memory. Imaging Neuroscience 2024;2:1–13. 10.1162/imag_a_00105.

[27] Clark IA, Kim M, Maguire EA. Verbal Paired Associates and the Hippocampus: The Role of Scenes. J Cogn Neurosci 2018;30:1821–45. 10.1162/JOCN_A_01315.

[28] Marks DF. Visual imagery differences in the recall of pictures. British Journal of Psychology 1973. 10.1111/J.2044-8295.1973.TB01322.X.

[29] Monzel M, Handlogten J, Reuter M. No verbal overshadowing in aphantasia: The role of visual imagery for the verbal overshadowing effect. Cognition 2024;245. 10.1016/J.COGNITION.2024.105732.

[30] Schmitt M, Altstötter-Gleich C, Hinz A, Maes J, Brähler E. Normwerte für das Vereinfachte Beck-Depressions-Inventar(BDI-V) in der Allgemeinbevölkerung. Diagnostica 2006;52:51–9. 10.1026/0012-1924.52.2.51/ASSET/IMAGES/LARGE/DIA5202051TBL5A.JPEG.

[31] Brenner D, Stirnberg R, Pracht ED, Stöcker T. Two-dimensional accelerated MP-RAGE imaging with flexible linear reordering. Magnetic Resonance Materials in Physics, Biology and Medicine 2014;27:455–62. 10.1007/S10334-014-0430-Y/TABLES/2.

[32] van der Kouwe AJW, Benner T, Salat DH, Fischl B. Brain morphometry with multiecho MPRAGE. Neuroimage 2008;40:559–69. 10.1016/J.NEUROIMAGE.2007.12.025.

[33] Stirnberg R, Stöcker T. Segmented K-space blipped-controlled aliasing in parallel imaging for high spatiotemporal resolution EPI. Magn Reson Med 2021;85:1540–51. 10.1002/MRM.28486.

[34] Stirnberg R, Huijbers W, Brenner D, Poser BA, Breteler M, Stöcker T. Rapid whole-brain resting-state fMRI at 3 T: Efficiency-optimized three-dimensional EPI versus repetition time-matched simultaneous-multi-slice EPI. Neuroimage 2017;163:81–92. 10.1016/J.NEUROIMAGE.2017.08.031.

[35] McIntosh AR. Mapping Cognition to the Brain Through Neural Interactions. Memory 1999;7:523–48. 10.1080/096582199387733.

[36] McIntosh AR, Lobaugh NJ. Partial least squares analysis of neuroimaging data: Applications and advances. Neuroimage, vol. 23, 2004. 10.1016/j.neuroimage.2004.07.020.

[37] Berk Mirza M, Adams RA, Mathys CD, Friston KJ. Scene construction, visual foraging, and active inference. Front Comput Neurosci 2016;10:196944. 10.3389/FNCOM.2016.00056/BIBTEX.

[38] Ladyka-Wojcik N, Liu ZX, Ryan JD. Unrestricted eye movements strengthen effective connectivity from hippocampal to oculomotor regions during scene construction. Neuroimage 2022;260. 10.1016/J.NEUROIMAGE.2022.119497.

[39] Zeidman P, Mullally SL, Maguire EA. Constructing, Perceiving, and Maintaining Scenes: Hippocampal Activity and Connectivity. Cerebral Cortex (New York, NY) 2015;25:3836. 10.1093/CERCOR/BHU266.

[40] Aly M, Ranganath C, Yonelinas AP. Detecting changes in scenes: The hippocampus is critical for strength-based perception. Neuron 2013;78:1127. 10.1016/J.NEURON.2013.04.018.

[41] Dalton MA, Zeidman P, McCormick C, Maguire EA. Differentiable processing of objects, associations, and scenes within the hippocampus. Journal of Neuroscience 2018;38:8146–59. 10.1523/JNEUROSCI.0263-18.2018.

[42] Lee SM, Shin J, Lee I. Significance of visual scene-based learning in the hippocampal systems across mammalian species. Hippocampus 2023;33:505–21. 10.1002/HIPO.23483.

[43] Lee ACH, Buckley MJ, Pegman SJ, Spiers H, Scahill VL, Gaffan D, et al. Specialization in the medial temporal lobe for processing of objects and scenes. Hippocampus 2005;15:782–97. 10.1002/HIPO.20101.

[44] Hodgetts CJ, Shine JP, Lawrence AD, Downing PE, Graham KS. Evidencing a place for the hippocampus within the core scene processing network. Hum Brain Mapp 2016;37:3779–94. 10.1002/HBM.23275.

[45] Pearson J. The human imagination: the cognitive neuroscience of visual mental imagery. Nat Rev Neurosci 2019. 10.1038/S41583-019-0202-9.

[46] Sheldon S, Levine B. The role of the hippocampus in memory and mental construction. Ann N Y Acad Sci 2016;1369:76–92. 10.1111/NYAS.13006.

[47] Robin J, Moscovitch M. Details, gist and schema: hippocampal–neocortical interactions underlying recent and remote episodic and spatial memory. Curr Opin Behav Sci 2017;17:114–23. 10.1016/J.COBEHA.2017.07.016.

[48] Monzel M, Leelaarporn P, Lutz T, Schultz J, Brunheim S, Reuter M, et al. Hippocampal-occipital connectivity reflects autobiographical memory deficits in aphantasia. Elife 2024;13. 10.7554/ELIFE.94916.1.

[49] Zvyagintsev M, Clemens B, Chechko N, Mathiak KA, Sack AT, Mathiak K. Brain networks underlying mental imagery of auditory and visual information. Eur J Neurosci 2013;37:1421–34. 10.1111/EJN.12140.

[50] De Luca F, McCormick C, Ciaramelli E, Maguire EA. Scene processing following damage to the ventromedial prefrontal cortex. Neuroreport 2019;30:828–33. 10.1097/WNR.0000000000001281.

[51] Kosslyn S, Thompson W. When is early visual cortex activated during visual mental imagery? Psychol Bull 2003. 10.1037/0033-2909.129.5.723.

[52] Kosslyn SM, Ganis G, Thompson WL. Neural foundations of imagery. Nat Rev Neurosci 2001. 10.1038/35090055.

[53] Klein I, Paradis AL, Poline JB, Kosslyn SM, Le Bihan D. Transient activity in the human calcarine cortex during visual-mental imagery: an event-related fMRI study. J Cogn Neurosci 2000;12 Suppl 2:15–23. 10.1162/089892900564037.

[54] Ishai A, Haxby J V., Ungerleider LG. Visual imagery of famous faces: Effects of memory and attention revealed by fMRI. Neuroimage 2002;17:1729–41. 10.1006/nimg.2002.1330.

[55] Lee S-H, Kravitz DJ, Baker CI. Disentangling visual imagery and perception of real-world objects. Neuroimage 2012. 10.1016/J.NEUROIMAGE.2011.10.055.

[56] Vannucci M, Pelagatti C, Chiorri C, Mazzoni G. Visual object imagery and autobiographical memory: Object Imagers are better at remembering their personal past. Memory 2016. 10.1080/09658211.2015.1018277.

[57] Cavanna AE, Trimble MR. The precuneus: a review of its functional anatomy and behavioural correlates. Brain 2006;129:564–83. 10.1093/BRAIN/AWL004.

[58] Spagna A, Hajhajate D, Liu J, Bartolomeo P. Visual mental imagery engages the left fusiform gyrus, but not the early visual cortex: A meta-analysis of neuroimaging evidence. Neurosci Biobehav Rev 2021;122:201–17. 10.1016/J.NEUBIOREV.2020.12.029.

[59] Conti F, Carnemolla S, Piguet O, Irish M. Scene construction in healthy aging – Exploring the interplay between task complexity and oculomotor behaviour. Brain Cogn 2024;177:106163. 10.1016/J.BANDC.2024.106163.

[60] Serre T, Crouzet SM, Linsley D, Roldan SM. Object Recognition in Mental Representations: Directions for Exploring Diagnostic Features through Visual Mental Imagery. Front Psychol 2017;8:833. 10.3389/FPSYG.2017.00833.

[61] Setton R, Wynn JS, Schacter DL. Peering into the future: Eye movements predict neural repetition effects during episodic simulation. Neuropsychologia 2024;197:108852. 10.1016/J.NEUROPSYCHOLOGIA.2024.108852.

